# Highly-efficient Cas9-mediated transcriptional programming

**DOI:** 10.1101/012880

**Authors:** Alejandro Chavez, Jonathan Scheiman, Suhani Vora, Benjamin W. Pruitt, Marcelle Tuttle, Eswar Iyer, Samira Kiani, Christopher D. Guzman, Daniel J. Wiegand, Dimtry Ter-Ovanesyan, Jonathan L. Braff, Noah Davidsohn, Ron Weiss, John Aach, James J. Collins, George M. Church

## Abstract

The RNA-guided bacterial nuclease Cas9 can be reengineered as a programmable transcription factor by a series of changes to the Cas9 protein in addition to the fusion of a transcriptional activation domain (AD)^1–5^. However, the modest levels of gene activation achieved by current Cas9 activators have limited their potential applications. Here we describe the development of an improved transcriptional regulator through the rational design of a tripartite activator, VP64-p65-Rta (VPR), fused to Cas9. We demonstrate its utility in activating expression of endogenous coding and non-coding genes, targeting several genes simultaneously and stimulating neuronal differentiation of induced pluripotent stem cells (iPSCs).

## RESULTS

Cas9 is an RNA-guided endonuclease that is directed to a specific DNA sequence through complementarity between the associated guide RNA (gRNA) and its target locus^6–9^. Cas9 can be directed to nearly any arbitrary sequence with a gRNA, requiring only a short protospacer adjacent motif (PAM) site proximal to the target^10–14^. Through mutational analysis, variants of Cas9 have been generated that lack endonucleolytic activity but retain the capacity to interact with DNA^1,9,15,16^. These nuclease-null or dCas9 variants have been subsequently functionalized with effector domains such as transcriptional activation domains (ADs), enabling Cas9 to serve as a tool for cellular programming at the transcriptional level^1–5^. The ability to programmably induce expression of a specific target within its native chromosomal context would provide a transformative tool for myriad applications, including the development of therapeutic interventions, genetic screening, activation of endogenous and synthetic genetic circuits, and the induction of cellular differentiation^17–20^.

To design an enhanced programmable activator, we screened a library of transcription factor ADs and members of the Mediator and RNA polymerase II complexes for potential use as transcriptional effectors^21^. More than 20 individual candidates were fused to the C-terminus of *Streptococcus pyogenes* (SP)-dCas9. These putative dCas9-activator constructs were transfected into mammalian cells and their potency assessed by quantifying fluorescence from a transcriptional reporter by flow cytometry^1^ (Supplemental Fig. 1). Of the hybrid proteins tested, three – dCas9-VP64, dCas9-p65 and dCas9-Rta – showed meaningful reporter induction^2,22^. Nonetheless, neither the p65 nor the Rta hybrid were stronger activators than the commonly used dCas9-VP64 protein (Supplemental Fig. 2).

In natural systems, transcriptional initiation occurs through the coordinated recruitment of the necessary machinery by a group of locally concentrated transcription factor ADs^23^. Hence, we hypothesized that joining multiple activation domains to a single dCas9 molecule would result in an increase in transcriptional activation by mimicking the natural cooperative recruitment process. Taking dCas9-VP64 as a starting scaffold, we introduced an additional nuclear localization signal to ensure efficient targeting. We subsequently extended the C-terminal fusion with the addition of either the p65 or Rta AD. As predicted, when p65 or Rta was joined to dCas9-VP64, we observed an increase in transcriptional output. Further improvement was observed when both p65 and Rta were fused in tandem to VP64, generating a hybrid VP64-p65-Rta tripartite activator (hereon referred to as VPR) (Fig. 1).

**Figure 1.**
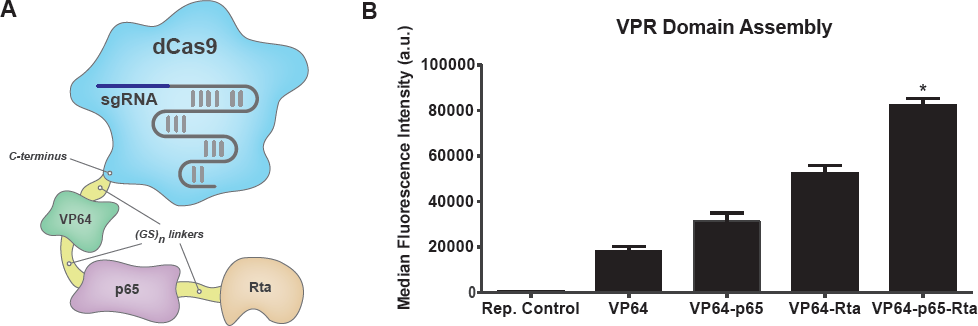
Activation is improved by serial fusion of activation domains to dCas9. (A) Transcriptional activation via Cas9 was performed by fusing three activation domains, VP64, p65, and Rta, to the C-terminus of a nuclease-null dCas9 protein. (B) Serial activation domain assemblies fused to dCas9 were tested against a fluorescent reporter assay. Data represent median fluorescence ± s.e.m. n=5 biological replicates, performed in a single experiment. (*denotes significance of dCas9-VP64-p65-Rta over all constructs including Reporter Control P = <0.0001, dCas9-VP64 P = <0.0001, dCas9-VP64-p65 P = <0.0001, and dCas9-VP64-Rta P = <0.0001).

To begin characterizing VPR, we confirmed the importance of each of its constituent parts (VP64, p65, and Rta) by replacing each member with mCherry and measuring the resulting protein’s activity by reporter assay. All fusions containing mCherry showed a decrease in activity relative to VPR but exhibited higher levels of activity with respect to the VP64 activation domain alone (Supplemental Fig. 3), demonstrating the essentiality of all members of the VPR complex. While the VPR fusion showed a clear improvement in transcriptional activation, the optimal ordering of the individual ADs remained unknown. To test the role of domain order, we designed a set of dCas9-VPR constructs in which we shuffled the positions of VP64, p65, and Rta to generate all possible non-repeating arrangements. Evaluation of the VPR permutations confirmed that the original VP64-p65-Rta order was indeed the optimal configuration (Supplemental Fig. 4).

Given the activation efficiency of our SP-dCas9-VPR fusion, we investigated whether the VPR construct would exhibit similar potency when fused to other DNA-binding scaffolds. Fusion of VPR to a nuclease-null *Streptococcus thermophilus* (ST1)-dCas9, a designer transcription activator like effector (TALE), or a zinc-finger protein allowed for a respective ∼6x, ∼4x, and ∼2x increase in activation relative to VP64, as determined by reporter assays (Supplemental Fig. 5)^19,24^.

Having performed initial characterization of our SP-dCas9-VPR fusion, we sought to assess its ability to activate endogenous coding and non-coding targets relative to VP64. To this end, we constructed 3-4 gRNAs against a set of factors related to cellular reprogramming, development, and gene therapy. When compared to the dCas9-VP64 activator, dCas9-VPR showed 22-to-320 fold improved activation of endogenous targets. Induced genes include the long-noncoding RNA (lncRNA) *MIAT*, the largest protein-coding gene in the human genome, *TTN*, several transcription factors, *NEUROD1*, *ASCL1*, *RHOXF2*, and a structural protein, *ACTC1* (Fig. 2A).

Cas9 enables multiplexed activation through the simple introduction of a collection of guide RNAs against a desired set of genes. To determine the efficiency of multi-gene targeting, we performed a pooled activation experiment simultaneously inducing four of our initially characterized genes *MIAT*, *NEUROD1*, *ASCL1*, and *RHOXF2*. VPR allowed for robust multi-locus activation, exhibiting several-fold higher expression levels than VP64 across the panel of genes (Fig. 2B).

The ability to regulate gene expression levels through transcriptional activation provides a powerful means to reprogram cellular identity for regenerative medicine and basic research purposes. Previous work has shown that the ectopic expression of several cDNAs enables cellular reprogramming of terminally differentiated cells to a pluripotent state, and can similarly induce differentiation of stem cells into multiple cell types^25^. While such studies typically require multiple factors, it was recently shown that exogenous expression of single transcription factors, Neurogenin2 (*NGN2*) or Neurogenic differentiation factor 1 (*NEUROD1*), is sufficient to induce differentiation of human iPS cells into induced neurons (iNeurons)^26,27^. Our group had previously attempted to recapitulate this same differentiation paradigm using dCas9-VP64 based activators and observed minimal differentiation activity (data not shown). We hypothesized this was due to insufficient activation and therefore postulated that VPR might overcome this barrier and enable differentiation of iPS cells to iNeurons by more potently activating either *NGN2* or *NEUROD1*.

**Figure 2.**
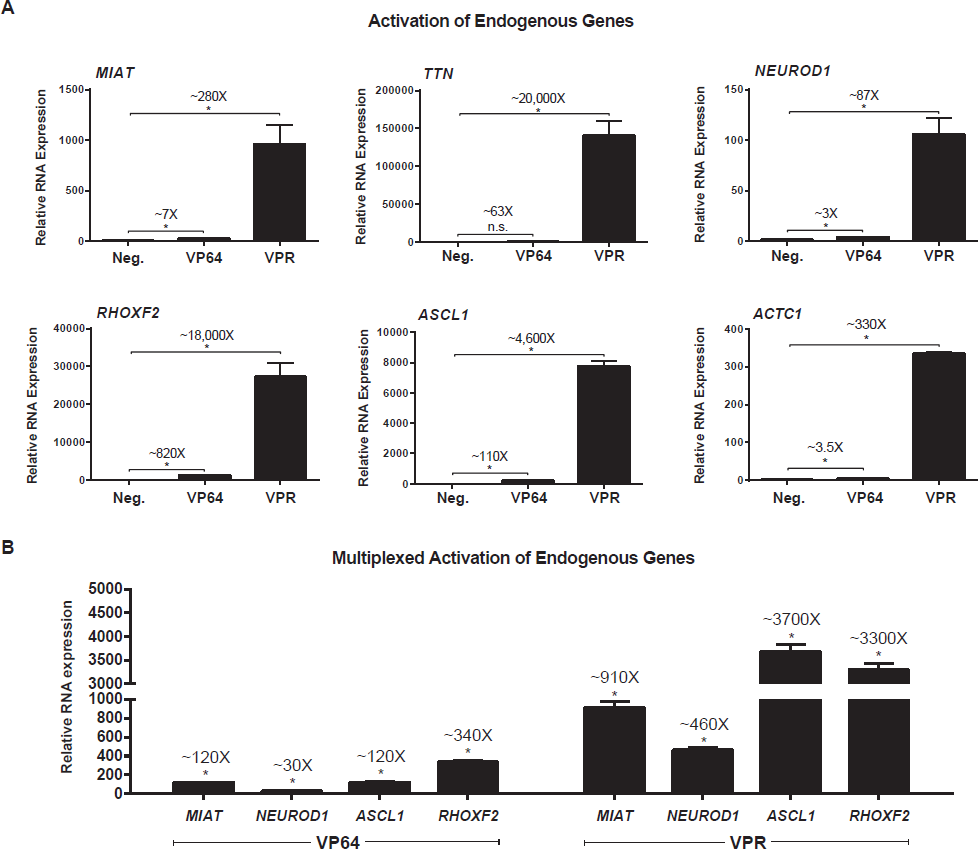
VPR represents a robust tool for gene activation. (A) Target gene expression, measured by qRT-PCR and normalized to Beta-actin mRNA levels, in HEK293T cells simultaneously transfected with 3-4 gRNAs targeting the indicated genes along with the labeled dCas9-activator construct (gRNAs used for activation are listed in the Supplemental Data). Negative controls were transfected with indicated gRNAs alone. Data are shown as the mean ± s.e.m (n = 3 biological replicates, performed in a single experiment). Difference in activation between VPR and VP64 was 40-fold, 320-fold, 26-fold, 22-fold, 40-fold, and 94-fold, for *MIAT*, *TTN*, *NEUROD1*, *RHOXF2*, *ASCL1*, and *ACTC1*, respectively. P values determined by two-tailed t-test (*dCas9-VP64 or dCas9-VPR versus gRNA-only control, respectively, *MIAT* P = 0.0277, 0.001, *TTN* P = n.s., 0.0003, *NEUROD1* P = 0.0009, 0.0003, *ASCL1* P = < 0.0001, <0.0001, *RHOXF2* P = < 0.0001, 0.0002, *ACTC1* P = 0.0002, <0.0001. n.s. = not significant. Comparison of dCas9-VP64 vs. dCas9-VPR, for all genes, is significant P = <0.0011). (B) Performed as in panel A except 3-4 gRNAs against each of the four endogenous genes, *MIAT*, *NEUROD1*, *ASCL1*, and *RHOXF2*, were transfected in unison. Data are shown as the mean ± s.e.m (n = 3 biological replicates, performed in a single experiment). P values determined by two-tailed t-test (*multiplexed dCas9-VP64 or dCas9-VPR vs. mock control sample for each gene respectively, *MIAT* P = 1.26 × 10^−5^, P = 1.26 × 10^−4^, *NEUROD1* P = 7.28 × 10^−5^, P = 4.72 × 10^−5^, *ASCL1* P = 5.82 × 10^−4^, P = 1.50 × 10^−5^, *RHOXF2* P = 1.57 × 10^−5^, P = 1.84 × 10^−5^. Comparison of dCas9-VP64 vs. dCas9-VPR, for all genes, is significant P = <0.0022).

Stable PGP1 iPS doxycycline-inducible dCas9-VP64 and dCas9-VPR cell lines were generated and transduced with lentiviral vectors containing a mixed pool of 30 gRNAs directed against either *NGN2* or *NEUROD1*. To determine differentiation efficiency, gRNA containing dCas9-AD iPS cell lines were cultured in the presence of doxycycline for four days and monitored for phenotypic changes (Supplemental Fig. 6 & 7). We observed that VPR, in contrast to VP64, enabled rapid and robust differentiation of iPS cells into a neuronal phenotype consistent with previously published reports. Additionally, these cells stained positively for the neuronal markers Beta III tubulin and neurofilament 200 (Fig. 3A and Supplemental Fig. 8A, respectively)^28^. Quantification of Beta III tubulin staining revealed that dCas9-VPR cell lines showed either ∼22.5× or ∼10× improvement in the amount of iNeurons observed through activation of *NGN2* or *NEUROD1*, respectively (Fig. 3B). Similar results were observed with neurofilament 200 staining (Supplemental Fig. 8B). Analysis by qRT-PCR four days after doxycycline-induction revealed a ∼10-fold and ∼18-fold increase in mRNA expression levels for *NGN2* and *NEUROD1*, respectively, within dCas9-VPR cells over their dCas9-VP64 counterparts (Fig. 3C).

**Figure 3.**
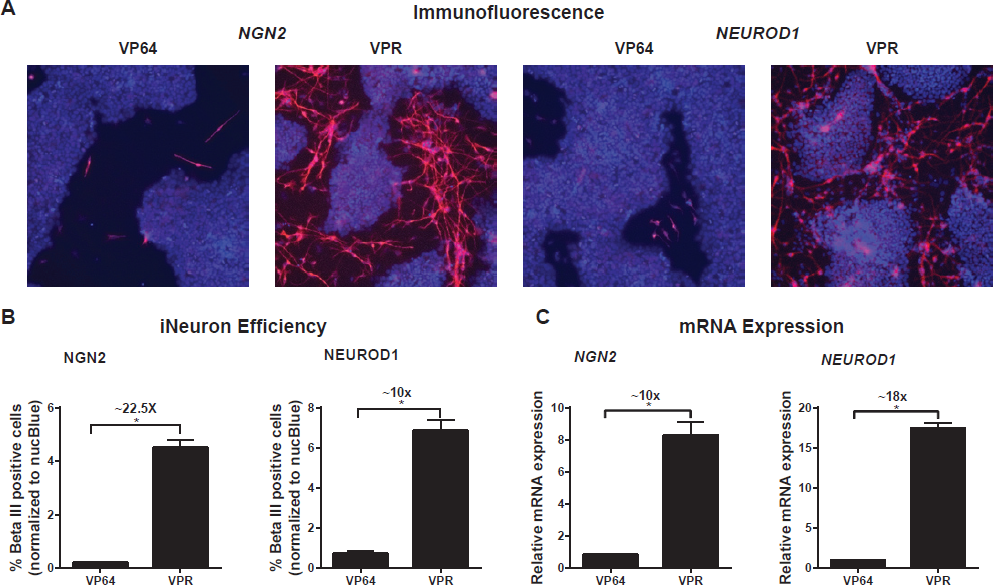
VPR is essential for efficient dCas9-mediated iPSC neuronal differentiation. (A) Immunofluorescence images for NucBlue (blue, total cells) and Beta III tubulin (red, iNeurons). Images were taken 4 days after doxycycline induction and are representative of biological triplicates. (B) Quantification and comparison of iNeurons generated by either dCas9-VP64 or dCas9-VPR in the presence of doxycycline. Percentages of iNeurons were determined by normalizing Beta III positive cells to total (NucBlue) cells. Data are shown as the mean ± s.e.m. (n=3 biological replicates, performed in a single experiment, with each replicate being an average of 24 separate images). P values determined by two-tailed t-test (*doxycycline induced dCas9-VPR versus doxycycline induced dCas9-VP64. P = 0.0002 and 0.0004 for *NGN2* and *NEUROD1*, respectively) (C) qRT-PCR analysis for mRNA expression levels of *NGN2* and *NEUROD1* in dCas9-AD iPS cell lines. Data is normalized to dCas9-VP64 cells and shown as the mean ± s.e.m. (n=2 biological replicates, performed in a single experiment). P values determined by two-tailed t-test (*doxycycline induced dCas9-VPR versus doxycycline induced dCas9-VP64. P = 0.0013 and <0.0001 for *NGN2* and *NEUROD1*, respectively).

## DISCUSSION

Here we introduce a highly-efficient Cas9 transcriptional activator. We demonstrate its versatility in both single and multi-target contexts along with its suitability for cellular reprogramming.

To identify more potent transcriptional effectors, we screened over 20 candidates previously shown to be involved in transcription initiation. While the majority of domains failed to show appreciable activity, the fusions with VP64 (derived from a herpes virus transcription factor), p65 (a member of the NFκB signaling pathway), and Rta (an activator of Epstein-Barr virus genes), showed appreciable activity when fused to dCas9. Previous biochemical analysis of these ADs has determined that each interacts with several members of the transcriptional and chromatin remodeling machinery – VP64 with MED17 and MED25, p65 with MED17, CBP, and P300, and Rta with TBP and TFIIB^21,29,30^. We speculate that the ability of VPR to simultaneously interact with multiple proteins involved in transcription explains the improvement in gene activation that we observe. Furthermore, given the abundance of factors with characterized roles in localized transcriptional initiation, we anticipate that further screening and combinatorial assembly of additional effector peptides will prove fruitful.

While VPR is a single tripartite activator construct fused to dCas9, recent evidence has demonstrated that multimeric recruitment of even the modestly effective VP64 AD can lead to abundant increases in transcriptional activation^31^. We expect that our more potent VPR fusion will be amenable to such multimerization strategies and thereby further enhance our ability to activate endogenous gene targets.

In this study, we focused primarily on identifying and engineering transcriptional ADs. Future designs, however, may directly employ a variety of regulatory proteins that alter chromatin state at the nucleotide, histone, and chromosomal levels^32^. This may prove useful for gaining access to target loci in heterochromatinized regions of the genome. Interestingly, preliminary evidence suggests that chromatin state may present a less significant impediment to programmable activation than anticipated; we were able to robustly activate *RHOXF2*, despite it being located in locus repressed by CpG methylation^33^. Therefore, VPR appears to be able to overcome some recalcitrant chromosomal states.

Previous work pioneered Cas9-mediated cellular differentiation, yet to our knowledge we are the first to use Cas9 effectors to successfully generate a targeted cell population (iNeurons)^34^. Furthermore, we demonstrate that effective differentiation is contingent upon the use of the more potent VPR activator. While the amount of iNeurons we observe upon dCas9-VPR mediated differentiation is less than 10% of the total cell population, it should be noted that the true proportion of differentiated cells is likely higher. Undifferentiated iPS cells continue to divide rapidly during our 4-day induction period while iNeurons are post-mitotic and thus remain under-represented at our analysis end-point^26,27^.

Another promising facet of our work is the ability to upregulate both endogenous coding and non-coding genes, paving the way for large-scale genomic screens free from constraints such as gene size or availability of previously constructed cDNA libraries^35^. This is particularly relevant in the context of lncRNAs, for which no large-scale expression library exists. Furthermore, by allowing for robust expression of genes from their native loci, we expect to maintain isoform heterogeneity within the particular cell type being tested, enabling a more thorough understanding of gene activity^36^. Finally, the ability to potently upregulate several target genes in unison will prove useful for future applications in directed cellular differentiation, genetic screening, gene network analysis and synthetic circuit engineering.

## MATERIALS AND METHODS

### Vectors used and designed

Activation domains were cloned using a combination of Gibson and Gateway assembly or Golden Gate assembly methods. For experiments involving multiple activation domains, ADs were separated by short glycine-serine linkers. All SP-dCas9 plasmids were based on Cas9m4-VP64 (Addgene #47319), ST1-dCas9 plasmids were based on M-ST1n-VP64 (Addgene #48675). Sequences for gRNAs are listed in the Supplemental Information. gRNAs for endogenous gene activation were selected to bind anywhere between 1 and 1000 bp upstream of the transcriptional start site. gRNAs for iPSC differentiation to iNeurons, targeting *NGN2* and *ND1*, were selected to bind anywhere between 1 and 2000 base pairs upstream of the transcriptional start site. All gRNAs were expressed from either cloned plasmids (Addgene #41817) or integrated into the genome through lentiviral delivery (plasmid SB700). gRNA sequences are listed within the Supplemental Data. Reporter targeting gRNAs were previously described (Addgene #48671 and #48672).

### Cell culture and transfections

HEK 293T cells were maintained in Dulbecco’s Modified Eagle’s Medium (Invitrogen) with high glucose supplemented with 10% FBS (Invitrogen) and penicillin/streptomycin (Invitrogen). Cells were maintained at 37°C and 5% CO_2_ in a humidified incubator. Cells were transfected in 24-well plates seeded with 50,000 cells per well, 200ng of dCas9 activator, 10ng of gRNA plasmid and 60ng of reporter plasmid (when required) were delivered to each well using 2.5 ul of Lipofectamine 2000. For multiplex activation 10ng of gRNA plasmid was used against each target (for example if three guide RNAs were used against a target, a total of 30ng of guide RNA plasmid was added). Cells were grown an additional 36-48 hours before being assayed using fluorescence microscopy, flow cytometry or lysed for RNA purification and quantification.

### Fluorescence Reporter Assay

SP-Cas9 reporter assays were performed by targeting all dCas9-ADs with a single guide to a minimal CMV promoter, driving expression of a fluorescent reporter. Addgene plasmid #47320 was used to screen for novel ADs (Supplemental Figures 1B and 2) or was altered to contain a sfGFP reporter gene instead of tdTomato for Figure 1B, Supplemental Figures 3 and 4. In addition, to control for transfection efficiency in Figure 1B, Supplemental Figures 3 and 4, an EBFP2 expressing control plasmid was co-transfected at 25ng per well (EBFP2 plasmid was not co-transformed in Supplemental Figure 2). To remove untransfected cells from the analysis, sfGFP fluorescence was only analyzed in cells with >10^3^ EBFP2 expression (as determined by flow cytometry). For fusion of VPR to other programmable transcription factors (ST1-dCas9, TALE, and zinc-finger protein) no EBFP2 plasmid was co-transfected. ST1-Cas9 reporter assays were performed using the previously described tdTomato reporter with an appropriate PAM inserted upstream of the Tdtomato coding region. (Addgene #48678)^24^.

### qRT-PCR analysis

RNA was extracted from cells using the RNeasy PLUS mini kit (Qiagen) according to manufacturer’s protocol. 500ng of RNA was used with the iScript cDNA synthesis Kit (BioRad), and 0.5ul of cDNA was used for each qPCR reaction, utilizing the KAPA SYBR® FAST Universal 2X qPCR Master Mix. qPCR primers are listed within the Supplemental Data.

### Lentivirus production

Lentiviral particles were generated by transfecting 293T cells with the pSB700 sgRNA expression plasmid (with cerulean reporter) and the psPAX2 and pMD2.G (Addgene #12260 and #12259) packaging vectors at a ratio of 4:3:1, respectively. Viral supernatants were collected 48-72h following transfection and concentrated using the PEG Virus Precipitation Kit (BioVision) according to the manufacturer’s protocol.

### iPSC culture and dCas9-AD cell line generation

PGP1 iPS cells were obtained from the Coriell Institute Biorepository (GM23338) and maintained on matrigel (Corning) coated tissue culture plates in mTeSR Basal medium (Stemcell technologies). To generate stable iPS Cas9-AD expressing cell lines, approximately 5×10^∧^5 cells were nucleofected with 1.5ug of Cas9-AD piggy-bac expression vector and 340ng of transposase vector (System Biosciences) using the Amaxa P3 Primary Cell 4D-Nucleofector X Kit (Lonza), program CB-150. Following electroporation, cells were seeded onto 24-well matrigel-coated plates in the presence of 10uM ROCK inhibitor (R&D systems) and allowed to recover for two days before expanding to 6-well plates in the presence of 20ug/ml hygromycin to select for a mixed population of Cas9-AD integrant containing cells.

### iPSC transduction and neural induction

For neural induction experiments, iPS Cas9-AD cell lines were transduced with lentiviral preparations containing 30 sgRNAs, targeted against *NEUROD1* or *NGN2*, one day after seeding onto matrigel coated plates. Transduced cells were expanded and then sorted for the top 15% of cerulean positive cells (pSB700 gRNA expression). Once sorted, sgRNA containing Cas9-AD iPS cell lines were obtained and seeded in triplicates onto matrigel coated 24-well plates with mTeSR + 10uM ROCK inhibitor, either in the presence or absence of 1ug/ml of doxycycline. Fresh mTeSR medium + or − doxycycline was added every day for 4 days, at which cells were analyzed by immunofluorescence and harvested for qRT-PCR analysis.

### Immunostaining of Cas9 iNeurons

All steps for staining were performed at room temperature. Samples were washed once with PBS then fixed with 10% formalin (Electron Microscopy Sciences) for 20 min followed by permeabilization with 0.2% Triton X-100/PBS for 15 min. Samples were then blocked with 8% BSA for 30 min followed by staining with primary antibodies diluted into 4% BSA. Staining was performed for either 3h with anti-Beta III eFluor 660 conjugate (eBioscience) or 1h with anti-neurofilament 200 (Sigma), both at a 1:500 dilution. Samples were then washed 3 times, 5 minutes each, with 0.1% tween/PBS, followed by one wash with PBS. For neurofilament 200 staining a secondary donkey anti-rabbit Alexa Fluor 647 (Life Sciences) antibody was added at a 1:1000 dilution in 4% BSA for 1h. Samples were again washed as previously mentioned then stained with nucBlue [Hoechst 33342] (Life Sciences) for 5 min.

### Image acquisition and analysis of Cas9 iNeurons

24-well plates stained for NucBlue and neuronal markers, were imaged with a 10x objective on a Zeiss Axio Observer Z1 microscope. Zen Blue software (Zeiss) was used to program acquisition of 24 images per well. Total cell (NucBlue) and iNeuron (Beta III tubulin or neurofilament 200) counts were quantified for each image using custom Fiji and Matlab scripts and used to determine the % of iNeurons per well by the formula:

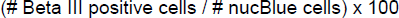

Images were post-processed and artificially colored for display in this publication (all adjustments were applied equally across the images).

### Statistical analysis

All statistical comparisons are two-tailed t-tests calculated using the GraphPad Prism software package (Version 6.0 for Windows. GraphPad Software, San Diego, CA). All sample numbers listed indicate the number of biological replicates employed in each experiment.

## ACKNOWLEDGEMENTS

We thank Kevin Esvelt, Prashant Mali and the rest of the members of the Church and Collins labs for helpful discussions. We thank Thomas C. Ferrante, Susan Byrne and Michael Farrell for technical assistance, Albert Keung for providing zinc-finger reporter constructs, and Joanna Schulak for graphic design. This work was supported by US National Institutes of Health NHGRI grant P50 HG005550, US Department of Energy grant DE-FG02-02ER63445 and the Wyss Institute for Biologically Inspired Engineering. A.C. acknowledges funding by the National Cancer Institute grant 5T32CA009216-34. S.V. acknowledges funding by the National Science Foundation Graduate Research Fellowship Program, the Department of Biological Engineering at MIT, and the Department of Genetics at Harvard Medical School.

## AUTHOR CONTRIBUTIONS

A.C. and J.R.S. conceived of the study. A.C., J.R.S., and S.V. designed and performed experiments and interpreted data. B.P. designed and developed fusion libraries. M.T., D.T-O., C.D.G., D.J.W., performed experiments. N.D. and J.L.B. developed reagents. E.I. performed the iNeuron analysis. S.K. and R.W. designed, tested and analyzed the TALE activator data. J.A. designed a subset of the gRNAs. J.J.C and G.M.C. supervised the study. A.C., J.R.S., S.V. and B.P. wrote the manuscript with support from M.T. and all authors.

